# *Cis*-regulatory architecture downstream of *FLOWERING LOCUS T* underlies quantitative control of flowering

**DOI:** 10.1101/2025.09.23.678055

**Authors:** Hao-Ran Zhou, Duong Thi Hai Doan, Thomas Hartwig, Franziska Turck

## Abstract

The *FLOWERING LOCUS T* (*FT*) gene is a central integrator of floral induction in *Arabidopsis thaliana*, with expression tightly regulated by complex transcriptional networks. Using CRISPR/Cas9 genome editing, we dissected the functional architecture of the *FT* downstream region and reveal that a 2.3-kb region immediately downstream of the *FT* coding sequence containing the Block E enhancer is essential for proper *FT* expression and flowering. Fine-scale deletions revealed a 63-bp core module with adjacent CCAAT- and G-boxes, whereas other conserved motifs had minor, context-dependent effects. We also uncovered a cryptic CCAAT-box module that becomes active when repositioned, coinciding with increased transcription factor binding and local chromatin accessibility, indicating that enhancer function is governed by local chromatin and motif context. The *cis*-regulatory logic revealed here provides insights into manipulating gene expression through the architecture and spatial arrangement of enhancer elements, potentially applicable beyond flowering genes or plant species.

## Main

Flowering time in Arabidopsis is triggered by the mobile florigen encoded by *FLOWERING LOCUS T* (*FT*), whose transcription is restricted to phloem companion cells of leaves before flowering and tightly tuned by environmental conditions and developmental stage^1–5^. Under long days (LD), *FT* transcripts peak at dusk, and in natural conditions show a bimodal pattern (morning and dusk), underscoring strong temporal control^6,7^. Understanding how *FT* integrates external with developmental cues is therefore central to explaining seasonal flowering.

Multiple flowering pathways converge at the *FT* locus through activators and repressors acting on defined *cis*-elements^8–10^. The key activator, CONSTANS (CO), tightly controlled by the circadian clock and light, promotes *FT* through CO-responsive elements (COREs) in its proximal promoter and acts phloem companion cells overlapping *FT*^3–5,11^. PHYTOCHROME INTERACTING FACTOR 4 (PIF4) and PIF7 activate *FT* under warm or shaded conditions by binding promoter and downstream sites^12–14^. MADS-box transcription factors, including FLOWERING LOCUS C (FLC), SHORT VEGETATIVE PHASE (SVP), and AGAMOUS-LIKE 15 (AGL15), repress *FT* via CArG-box motifs in the promoter and first intron, with AGL15 additionally binding downstream^15–17^. AP2-like repressors, such as SCHLAFMÜTZE (SMZ) and TARGET OF EAT1 (TOE1), suppress *FT* either directly, by binding promoter and downstream regulatory regions^18–20^, or indirectly, by interfering with CO function^21^. Together, these regulators act through *cis*-elements distributed across the *FT* locus to ensure tight spatiotemporal control.

Our previous work identified conserved promoter elements, including Blocks A, B, and C, that contribute strongly to *FT* regulation^4,22,23^. Beyond promoter regions, *cis*-regulatory elements can also occur downstream, where they function as transcriptional enhancers that integrate developmental and environmental cues^24–26^. Such downstream enhancers are increasingly recognized across eukaryotes, including plants, animals, and fungi, and can act over large genomic distances^27–31^. We recently identified Block E, an accessible region ∼1.4 kb downstream of *FT*, as a novel enhancer^32^. Block E is conserved across Brassicaceae and harbors multiple conserved transcription factor binding motifs: two CCAAT-boxes, a G-box, an I-box, a RE-α, and a TBS-like motif^20,23,32^. It overlaps with binding sites of both activators (PIF4, PIF7) and repressors (AGL15, SMZ, TOE1)^17–20,33,34^. The CCAAT-box is one of the most frequently occurring *cis*-elements in eukaryotic promoters and enhancers, recognized by the conserved NF-Y heterotrimeric transcription factor complex^35–37^. The G-box is probably the direct binding site for activators PIF4 and PIF7, while the TBS-like motif is bound by AP2-like repressors TOE1^20^. Yet it remains unclear whether additional essential elements lie downstream, which Block E motifs are required *in vivo*, and whether Block E acts as a multifunctional platform that integrates both activating and repressive signals.

Here, using CRISPR/Cas9 editing at the native locus, we show that a 2.3-kb region immediately downstream of *FT* (including Block E) is indispensable for *FT* expression and flowering, whereas more distal sequences are largely dispensable. Fine dissection identifies a 63-bp core in Block E, comprising a unit of one closely spaced CCAAT- and G-box, as necessary for enhancer activity; other motifs contribute modestly and context-dependently. We also uncover a latent downstream CCAAT-box module that becomes active when repositioned closer to Block E. Our findings reveal a modular, position-sensitive downstream architecture of *cis*-regulatory code that confers robustness and plasticity to *FT* regulation.

## Results

### Identification of potential regulatory elements including expressed long non-coding RNAs downstream of *FT*

To investigate the presence of additional *cis*-regulatory elements in the downstream region, we analyzed a 14.3-kb sequence downstream of *FT* that contains only two small peptide-coding genes, *AT1G65481* and *AT1G65483* (Fig. 1a). Comparative genomic analysis of *A. thaliana* and five related species revealed high conservation in one coding gene (*AT1G65481*) and four non-coding regions: Block E, *AT1NC09061*, *AT1G08757*, and *AT1G08763* (Fig. 1a). *AT1G08757* and *AT1G08763* are annotated as long non-coding RNAs (lncRNAs) in Araport11, while *AT1NC09061*, although not annotated, was identified as a lncRNA in a large-scale tiling array analysis^38^. Despite most of the 14.3-kb region being enriched in H3K27me3, two prominent open chromatin regions were detected in leaf: one encompassing Block E and another at the transcription start site of *AT1G08757* (Fig. 1a). Given the potential involvement of lncRNAs in gene regulation, these three conserved lncRNAs (*AT1NC09061*, *AT1G08757*, and *AT1G08763*) were considered candidate regulatory elements of *FT*.

**Fig. 1:**
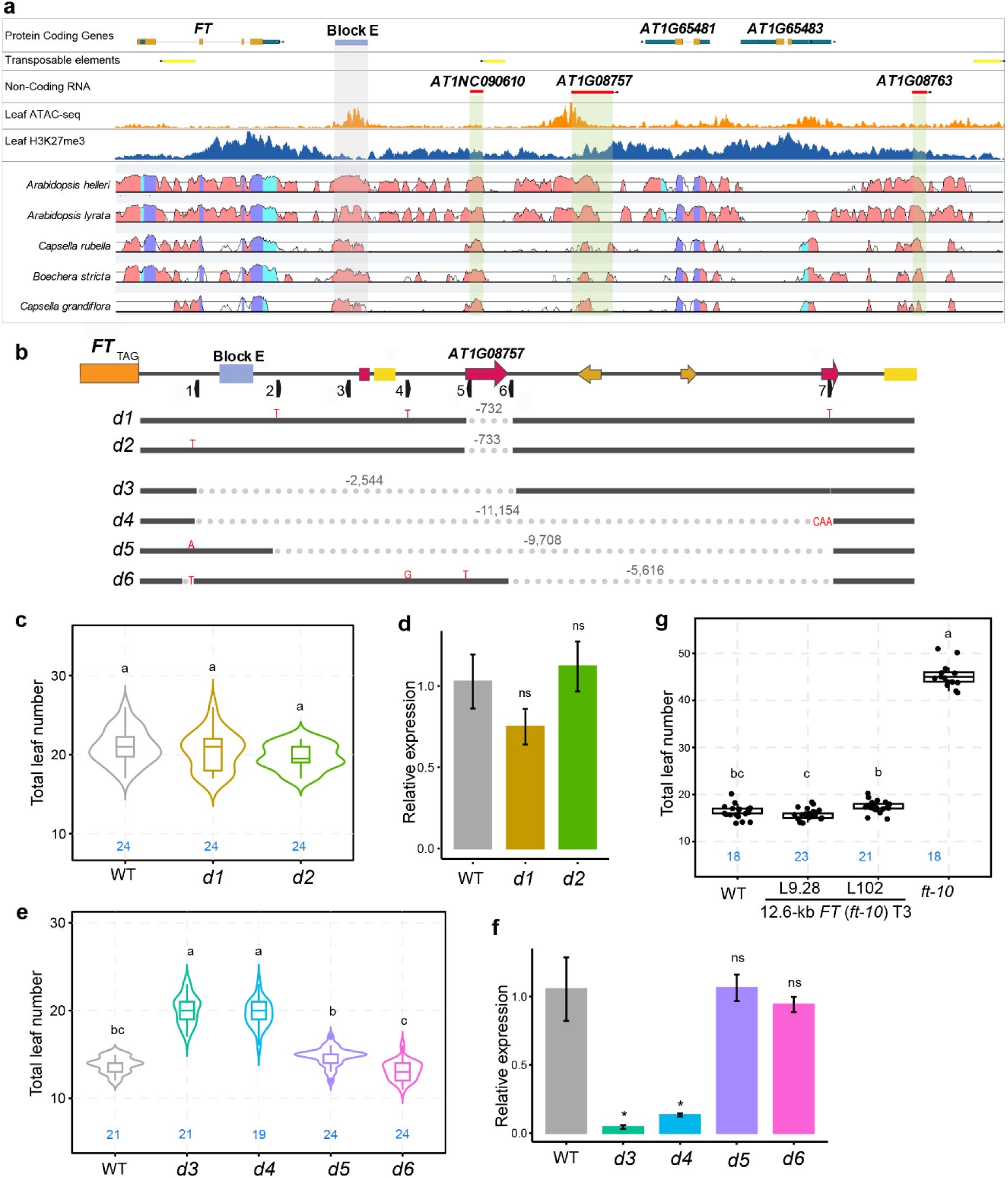
The immediate 2.3-Kb downstream region of *FT* is essential for proper expression and flowering. **a**, Jbrowse scheme of the *FT* locus and 14 Kb downstream region. Positions of Protein coding genes, transposable elements and non-coding RNAs as indicated as in the Araport11 annotation, the positions of the downstream *FT* enhancer Block E are indicated as blue box and gray shadow. Tracks ATAC and H3K27me3 indicate ATAC-seq and ChIP-seq coverage tracks for open chromatin and PcG targeted chromatin, respectively. Vista plot indicates percent identity to syntenic regions in other Brassicaceae. **b**, Scheme of 2.5 Kb *FT* down-stream region and nature of 6 deletion variants (*d1*-*d6*) generated by CRISPR/Cas9 genome editing. The locations of genomic features are indicated by boxes: Block E (blue), non-coding transcripts (red), transposable elements (yellow), transcripts annotated as coding (orange). **c**, Flowering time measured as leaves produced at the main shoot until flowering in Col-0 and two independent deletion mutants (*d1*, *d2*) of non-coding transcript *AT1G08757*. The number of replicates is indicated in blue, statistical test by ANOVA with posthoc Tukey HSD testing. Statistically significant differences are indicated by different letters. **d**, *FT* transcript accumulation measured by RT-qPCR for samples collected at ZT16 from 12-day old seedlings grown in LD. Analysis was using the 2deltaCq method using Col-0 as reference and *PP2A* as housekeeping control. Genotypes as in **c**. **e**, Analysis of flowering time as in **c** for two deletions encompassing Block E (*d3*, *d4*) and two deletions downstream of Block E (*d5*, *d6*). **f**, *FT* transcript accumulation measured as in **d** for genotypes used in **e**. **g**, Flowering time measured as in **c** of two independent single-copy transgenic lines containing a 12.6-Kb genomic region of *FT* that includes all regulatory regions from Chr1:24323415 to Chr1:24335986. Boxplots in c and e denote the range from the first to the third quartile, lines within boxes indicate the median and whiskers represent 1.5-fold of the interquartile range.

We then assessed the diurnal expression patterns of *FT* and these three lncRNAs in *Arabidopsis* Col-0 wild type grown under long-day (LD) conditions. As previously reported^7,39–41^, *FT* displayed a clear diurnal rhythm with a small morning peak and a larger dusk peak (Extended Data Fig. 1a). Expression of *AT1NC09061* was undetectable in our experiments, suggesting very low or silenced expression, possibly due to its proximity to a transposon (Fig. 1a). Interestingly, *AT1G08757* also exhibited a clear diurnal expression pattern, with a morning trough and a flattened evening peak (Extended Data Fig. 1b), whereas *AT1G08763* showed no discernible diurnal rhythm (Extended Data Fig. 1c).

The spatial expression pattern of *AT1G08757* was further examined using GUS staining (Extended Data Fig. 1d). Transgenic lines harboring the *AT1G08757p-GUS* construct showed GUS activity in the vasculature of cotyledons and the distal regions of the first true leaves, resembling the *FT* promoter-driven expression pattern, with additional signals at leaf hydathodes.

Together, these analyses show that three conserved lncRNAs reside within the 14.3-kb downstream region of *FT*, of which *AT1G08757* displays spatial and temporal expression patterns overlapping with *FT*.

### The immediate 2.3-Kb downstream region of *FT* is essential for its expression and floral transition

To assess the functional significance of the downstream lncRNAs and broader regulatory region, we generated CRISPR/Cas9 deletions across the 14.3-kb interval downstream of *FT*. Seven sgRNAs were assembled into a Cas9 expression vector and transformed into Col-0 (Fig. 1b). Deletion of the annotated *AT1G08757* (*d1*, *d2*) had no effect on flowering time or *FT* expression under LD conditions (Fig. 1c, d), indicating that *AT1G08757* is dispensable for *FT* regulation in these conditions.

We then generated larger deletions spanning 2.5–11 kb (*d3*–*d6*; Fig. 1b). Under LD conditions, only *d3* and *d4* mutants, which removed Block E, showed delayed flowering and reduced *FT* expression, whereas *d5* and *d6*, which retained Block E but eliminated downstream lncRNAs, were indistinguishable from wild type (Fig. 1e, f). These results imply Block E is essential for *FT* expression and floral induction, while excluding a functional role for the lncRNAs.

To further test the sufficiency of the downstream interval, we performed complementation assay using a 12.6-kb *FT* genomic fragment encompassing the 8.1-kb promoter, coding region, and 2.3-kb downstream sequence (Extended Data Fig. 2a). T2 populations of five independent lines segregated into wild-type–like and *ft-10*–like flowering, reflecting transgene presence (Extended Data Fig. 2b). Early flowering in L6 and L7 correlated with multiple transgene copies, whereas single-copy lines L8 and L9 flowered near WT, with L9 showing minimal variation (Extended Data Fig. 2b, c). Homozygous T3 plants from two independent lines flowered uniformly at WT timing under long-day conditions (Fig. 1g), demonstrating that a single copy of this fragment, including the 2.3-kb downstream region, is sufficient for LD-induced *FT* expression and flowering.

### A 63-bp sequence containing CCAAT-box within Block E is crucial for Block E‘s enhancer activity

To explore the function of the 2.3 kb downstream region of *FT*, we generated a series of mutants using CRISPR/Cas9 and guide RNAs targeting Block E and its surrounding sequences (Fig. 2a). Comparative sequence analysis across six Brassicaceae species revealed strong conservation of the I-box, the first CCAAT-box, and the RE-α motif (Extended Data Fig. 3). The G-box and TBS-like motifs were also highly conserved but contained a single substitution in the two Capsella species. The second CCAAT-box was restricted to Arabidopsis and Boechera, while the PBE-box was unique to *A. thaliana*. A fully conserved TATA-box further suggested that Block E carries promoter-like features.

**Fig. 2:**
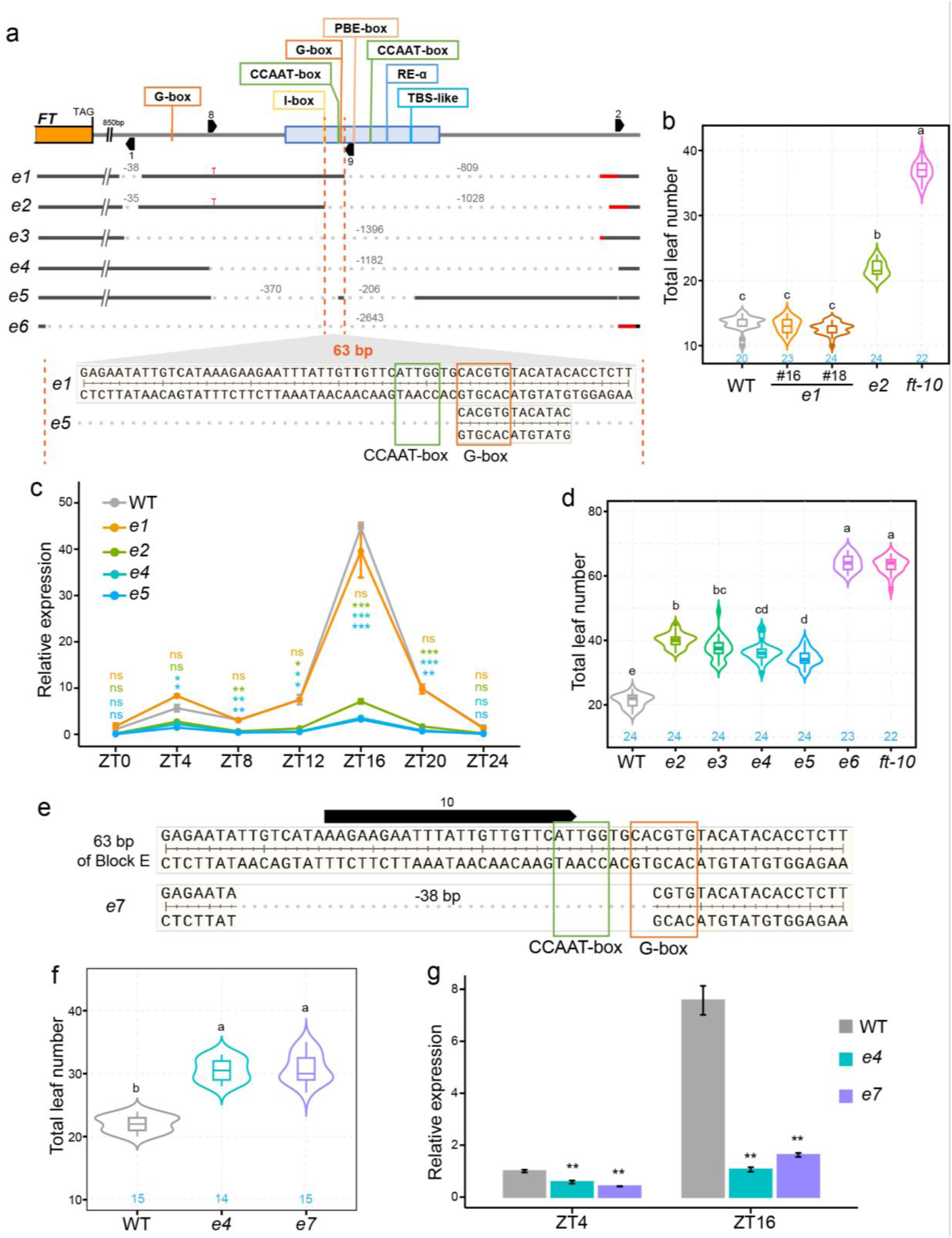
CCAAT-box and G-box are crucial for Block E in promoting *FT* expression and floral transition. **a**, Scheme of 2.5 Kb of *FT* down-stream region and nature of 6 Block E deletion variants (*e1*-*e6*) generated by CRISPR/Cas9 genome editing. The locations of Block E are indicated by a blue box, the locations of relevant *cis*-motifs are indicated by colored lines. The positions of sgRNAs used for genome editing are indicated by black arrowed boxes. **b**, Flowering time measured as leaves produced at the main shoot until flowering in wild type (WT), two independent segregants of *e1*, *e2*, and *ft-10* grown in the greenhouse. The number of replicates is indicated in blue, statistical test by ANOVA with posthoc HSD Tukey testing. Statistically significant differences are indicated by different letters. **c**, *FT* transcript accumulation from *e1*, *e2*, *e4*, *e5*, and WT measured by RT-qPCR for samples collected at 4h intervals during a period of 24h from 12-day old seedlings grown in LD. Analysis was using the 2deltaCq method using Col-0 as reference and *PP2A* as housekeeping control. Statistical analysis by Welch’s *t*-test, significant differences indicated by stars (ns p>0.05, * p<0.05, ** p<0.01, *** p<0.001). **d**, Flowering time analyzed as in **b** for deletions (*e2*-*e5*) that included a 63 bp region of Block E which contains a CCAAT-box and a G-box. Col-0 and *ft-10* were used as wild-type and loss-of-function control, deletion *e6* includes a partial deletion of the *FT* coding region. **e**, Detailed view of the 63 bp sequence of Block E in WT and the 38 bp deletion line *e7* that lack the CCAAT-box and the first two base pairs of the G-box. The position of sgRNA 10 is indicated above with a black arrowed box. **f**, Flowering time analyzed as in **b** for deletions *e4* and *e7*. **g**, *FT* transcript accumulation from genotypes as in **f** measured as in **d**.

Mutant *e1*, lacking the PBE-box, second CCAAT-box, RE-α, and TBS-like motifs, flowered slightly earlier than wild type in some experiments, or showed a weak early-flowering tendency in others (Fig. 2b and Extended Data Fig. 4a). *FT* expression analysis over a 24-hour time course showed a marginal, non-significant increase at ZT4 in *e1* seedlings (Fig. 2c), suggesting a role for these motifs in repression, potentially through AP2-like factors acting at the TBS-like site. In contrast, *e2*, which carries deletions similar to *e1* plus an additional 63-bp deletion encompassing the first CCAAT-box and G-box within Block E (Fig. 2a), flowered significantly later than wild type though earlier than *ft-10*, with strongly reduced *FT* expression across the day (Fig. 2b, c, d). The phenotypic contrast between *e1* and *e2* highlights the contribution of the 63-bp sequence to Block E enhancer activity.

Additional late-flowering Block E mutants (*e3*–*e5*) were identified. Deletion of the entire Block E (*e3*, *e4*) or almost all of it while retaining only 13 bp of the critical 63-bp region (*e5*) consistently reduced *FT* expression and delayed flowering, but did not alter its diurnal rhythm (Fig. 2c, d), indicating Block E controls expression amplitude rather than rhythmicity and the Block E G-box alone is insufficient for full enhancer activity. Additionally, the presence of G-box in immediately upstream of Block E did not compensate for enhancer loss, as *e2*, *e3*, and *e4* phenotypes were comparable (Fig. 2d). As a control, *e6*, disrupting the last exon of *FT* (Fig. 2a), was indistinguishable from *ft-10* (Fig. 2d).

Public PIF7 ChIP-seq data showed strong enrichment at Block E specifically under low red/far-red (R/FR) light^34^, consistent with shade-induced *FT* activation^13,14^ (Extended Data Fig. 4b, c). In agreement, *FT* expression was reduced in the *pif457* triple mutant only under far-red light (WL+FR), whereas Block E mutants *e3*, *e4*, and *e5* exhibited reduced *FT* expression under both control (WL) and far-red light conditions (Extended Data Fig. 4c). Although far-red light elevated *FT* expression across all genotypes, *FT* levels in *e3*, *e4*, and *e5* remained consistently lower than wild type, with no significant differences among the three mutants (Extended Data Fig. 4c). Similar to *pif457*, the induction ratio of *FT* by far-red light (WL+FR/WL) was significantly reduced in the Block E mutants compared to wild type (Extended Data Fig. 4c). Together, these results demonstrate Block E is required for maintaining *FT* expression under both normal and shade conditions.

To specifically test the importance of the first CCAAT-box and G-box, we generated mutant *e7* with a 38-bp deletion within the critical 63-bp region of Block E, disrupting both motifs (Fig. 2e). *e7* flowered as late as *e4* and had similarly reduced *FT* expression at ZT4 and ZT16 (Fig. 2f, g), confirming that both motifs contribute to Block E enhancer function, though their individual roles remain unresolved. Collectively, these results establish Block E as a critical enhancer of *FT* expression, with the 63-bp region containing the first CCAAT-box as central to its regulatory function.

### Minor contributions of the second CCAAT-box, RE-alpha, and TBS-like motifs in Block E to *FT* expression and flowering

To further dissect the contribution of individual transcription factor binding sites within Block E, we generated CRISPR/Cas9 mutants targeting the second CCAAT-box (*e8*), RE-alpha (*e9*), and TBS-like (*e10*) motifs. In *e8*, 27-bp encompassing the second CCAAT-box was replaced by 5 bp; In *e9*, a 67-bp fragment containing RE-alpha was deleted; and in *e10,* a 46-bp deletion removed the TBS-like motif (Fig. 3a). In contrast to the first CCAAT-box and G-box, which are essential for Block E enhancer function (Fig. 2), deletion of these individual sites had limited effects on flowering time and *FT* expression.

**Fig. 3:**
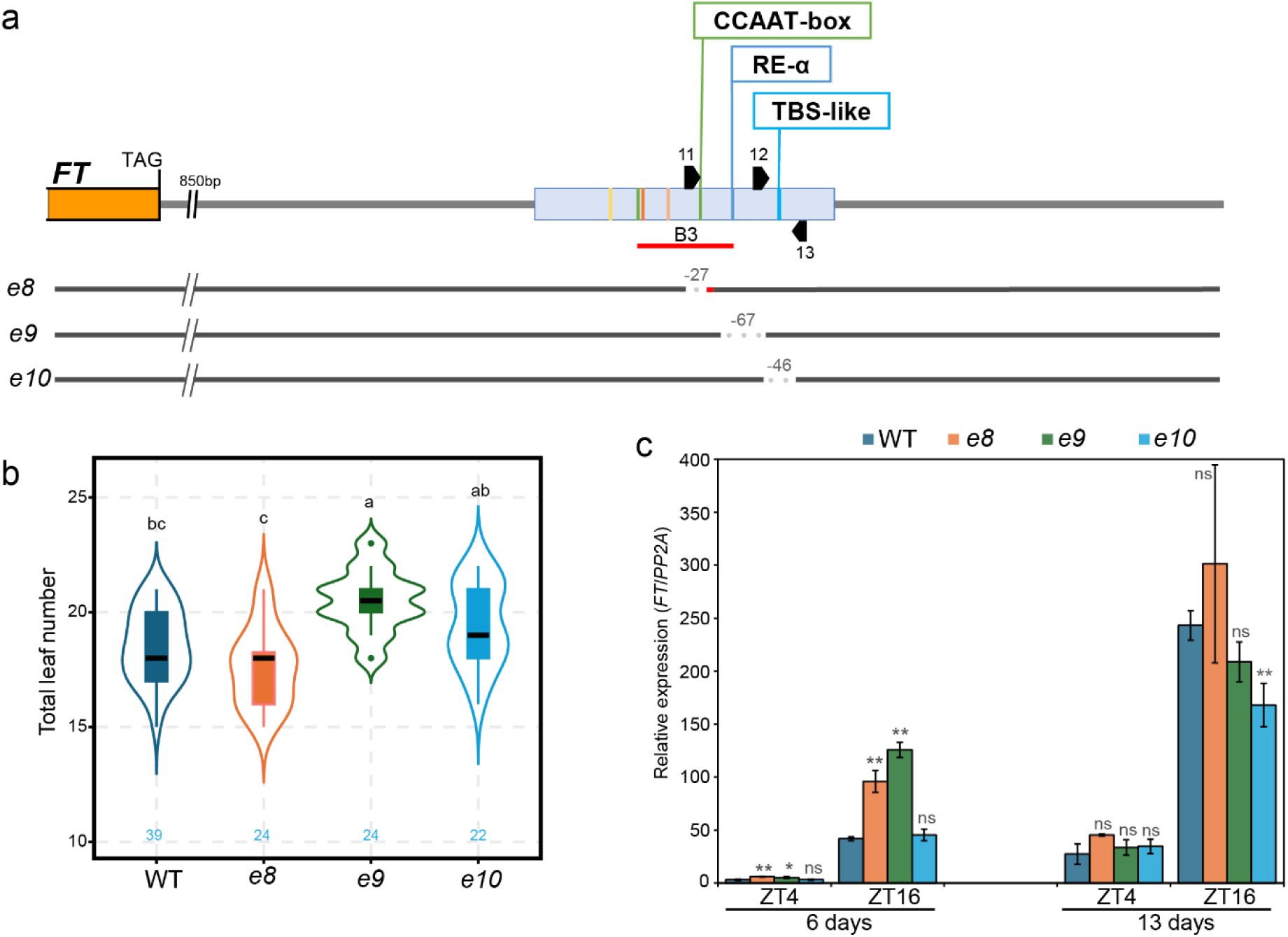
Deletion of the second CCAAT-box, RE-alpha, or TBS-like motifs in Block E has minor effects on *FT* and flowering. **a**, Schematic representation of the 2.5 kb downstream region of *FT* and the CRISPR/Cas9-generated Block E mutants *e8*, *e9*, and *e10*. Block E is indicated by a blue box. Colored lines within the blue box denote the positions of specific motifs, and the second CCAAT-box, RE-alpha, and TBS-like motifs are labeled above as named boxes. Black arrowheads indicate the positions of sgRNAs used for genome editing. The red underline marks the B3 region associated with AGL15 binding. **b**, Flowering time, measured as the total number of leaves produced at the main shoot until flowering, in wild type (WT) and the three mutants (*e8*, *e9*, and *e10*), each targeting a different motif within Block E. Statistical test by ANOVA with posthoc HSD Tukey testing. Statistically significant differences are indicated by different letters. **c**, *FT* transcript accumulation from genotypes as in **b** measured by RT-qPCR for samples collected at ZT4 and ZT16 from 6- and 13-day-old seedlings grown under long-day (LD) conditions. Analysis was using the 2deltaCq method using Col-0 as reference and *PP2A* as housekeeping control. Statistical analysis by Welch’s *t*-test, significant differences indicated by stars (ns p>0.05, * p<0.05, ** p<0.01, *** p<0.001).

Plants carrying the *e8* mutation showed a slight but non-significant trend toward earlier flowering compared to wild type (Fig. 3b). *FT* expression was consistently elevated in 6-day-old seedlings but indistinguishable from wild type by day 13. Notably, the deleted sequence overlaps with a previously defined AGL15-binding region B3 (Fig. 3a), suggesting a potential role in modulating MADS-domain protein enrichment at Block E.

By contrast, the *e9* and *e10* mutants showed delayed flowering relative to WT, with *e9* exhibiting a stronger delay and *e10* an intermediate phenotype (Fig. 3b). In *e9*, *FT* expression was elevated in 6-day-old seedlings, but in 13-day-old seedlings expression was similar to wild type at ZT4 and not significantly reduced at ZT16 (Fig. 3c). In *e10*, *FT* expression remained largely unchanged except for a significant reduction at ZT16 in 13-day-old seedlings (Fig. 3c).

These findings indicate that the second CCAAT-box, RE-alpha and TBS-like motifs each contribute modestly to Block E activity but are not individually required for robust *FT* induction or timely flowering under long-day conditions. This limited impact may reflect context-dependent or stage-specific roles for these motifs as indicated by the age-dependent changes in *FT* expression.

### A downstream CCAAT-box module may enhance or partially compensate for Block E activity

The extension of the *e1* to *e4* deletions beyond Block E prompted us to further investigate the regulatory role of the Block E downstream sequence. We used six CRISPR/Cas9 guide RNAs targeting both Block E and its immediate downstream region, generating mutants *e11 to e16* (Fig. 4a).

**Fig. 4:**
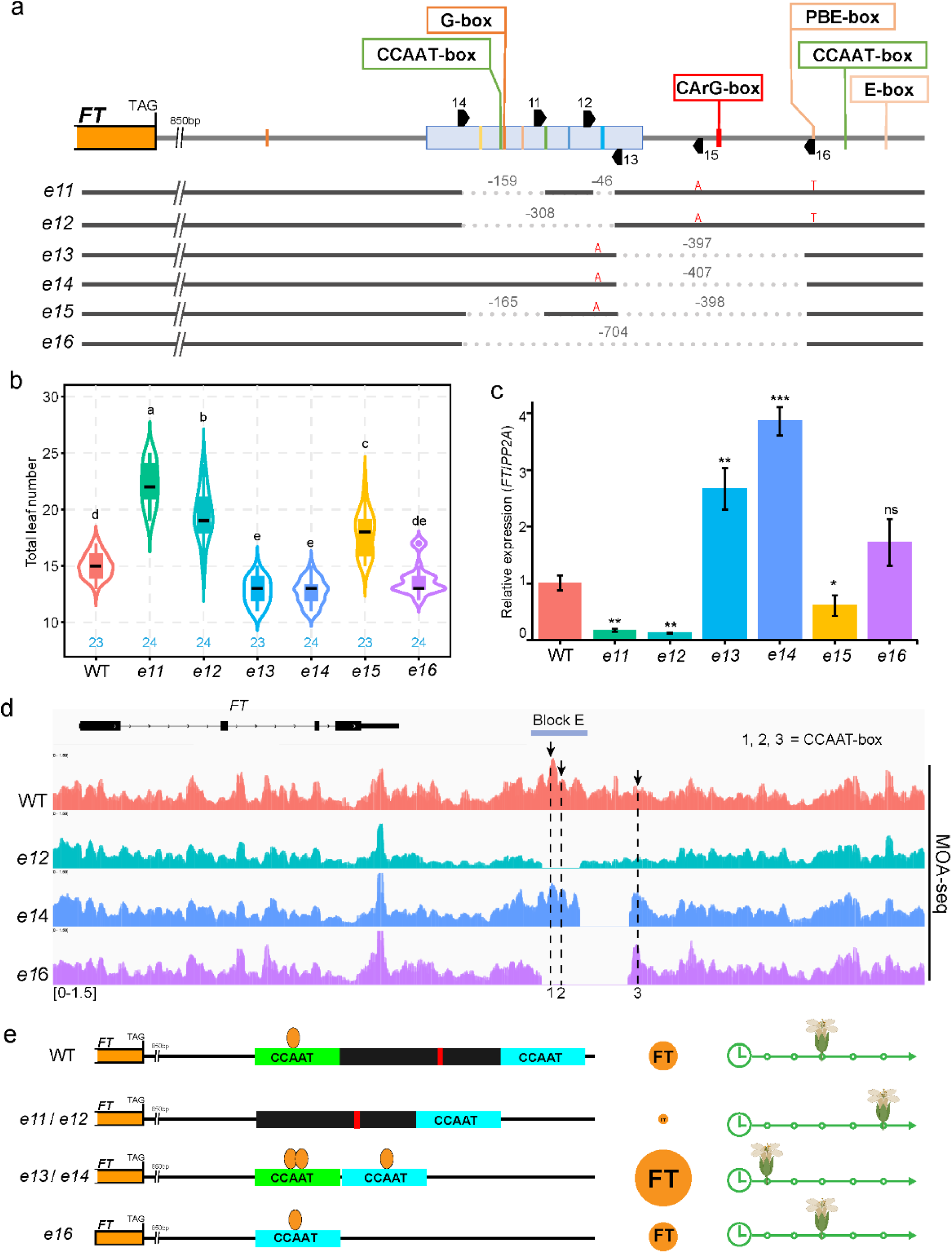
Regulatory impact of a latent CCAAT-box module downstream of Block E on floral transition. **a**, Schematic representation of the 2.5 kb *FT* downstream region and the nature of CRISPR/Cas9-generated deletion variants (*e11*–*e16*). Block E is indicated by a blue box. Relevant *cis*-regulatory motifs within and downstream of Block E are shown as colored lines, with selected motifs labeled above as named boxes. The positions of sgRNAs used for genome editing are marked with black arrowheads. **b**, Flowering time measured as leaves produced at the main shoot until flowering in Col-0 and deletion lines *e11*-*e16* as indicated in **a**. Statistical test by ANOVA with posthoc HSD Tukey testing. Statistically significant differences are indicated by different letters. **c**, *FT* transcript accumulation from genotypes as in **b** measured by RT-qPCR for samples collected at ZT16 from 12-day old seedlings grown under long-day (LD) conditions. Analysis was using the 2deltaCq method using Col-0 as reference and *PP2A* as housekeeping control. Statistical analysis by Welch’s *t*-test, significant differences indicated by stars (ns p>0.05, * p<0.05, ** p<0.01, *** p<0.001). **d**, Overlay of MOA-seq read coverage from two biological replicates at the *FT* locus and downstream region in wild type (red), *e12* (cyan), *e14* (blue), and *e16* (purple). The *FT* gene structure is shown above (black boxes, exons and UTRs; black lines, introns). Block E is indicated by a light blue line. Numbers 1–3 denote the three CCAAT-box motifs within Block E and its downstream region; their exact positions are indicated by black arrows and corresponding dashed lines. **e**, Model illustrating the positional and contextual effects of the downstream CCAAT-box on *FT* expression and flowering time. The downstream CCAAT-box, located beyond Block E, can compensate for the loss of the Block E CCAAT-box and enhance expression when repositioned adjacent to it. The green box labeled “CCAAT” marks the Block E module, and the blue box labeled “CCAAT” marks the downstream module. A black box represents a regulatory barrier, with the embedded red line indicating the CArG-box. Orange ovals represent transcription factors (e.g., NF-Y). Orange circles labeled “FT” indicate *FT* expression, with circle size reflecting relative expression levels. Clocks with arrows and flowers depict flowering time, with flower position along the arrow indicating earlier or later flowering.

Mutants *e11* and *e12* had deletions largely confined to Block E. All known Block E motifs (including the first CCAAT-box and G-box) were removed in both mutants, with only the second CCAAT-box and RE-alpha motif retained in *e11* (Fig. 4a). Consistent with the essential roles of the first CCAAT-box and G-box, both *e11* and *e12* exhibited delayed flowering and markedly reduced *FT* expression compared to wild type (Fig. 4b, c). Surprisingly, mutants *e13* and *e14*, which carried ∼400-bp deletions immediately downstream of Block E, flowered significantly earlier and displayed remarkably higher *FT* transcript levels than wild type (Fig. 4a, b, c). This early-flowering phenotype depended on Block E integrity, as *e15* (harboring the same downstream deletion as *e13*/*e14* but lacking the first CCAAT-box and G-box of Block E) flowered later than wild type, albeit earlier than *e11* and *e12*, and showed intermediate *FT* expression levels (Fig. 4a, b, c). Mutant *e16*, which deleted both Block E and the adjacent downstream region, had minimal impact on flowering or *FT* expression (Fig. 4a, b, c), suggesting a neutralization of the individual effects of Block E and the immediate downstream sequence when both are deleted.

These contrasting outcomes suggested that the downstream region normally exerts a repressive effect and that its removal alters the functional output of Block E. Sequence analysis revealed a repressive CArG-box motif within the downstream regions deleted in *e13* to *e16* (Fig. 4a). Intriguingly, a PBE-box, CCAAT-box, and E-box module resides immediately downstream of these deleted regions (Fig. 4a). We therefore hypothesized that loss of the CArG-box, coupled with repositioning of these downstream *cis*-elements, could reshape transcription factor binding at Block E and the downstream module, thereby enhancing or compensating *FT* activation.

To test this hypothesis, we performed Micrococcal Nuclease-defined cistrome Occupancy Analysis (MOA-seq) in wild type, *e12*, *e14*, and *e16* (Fig. 4d and Extended Data Fig. 5). In wild type, strong TF (likely NF-Y complexes) binding was detected at the first CCAAT-box within Block E, with weaker occupancy at the second CCAAT-box and no clear binding at the downstream (third) CCAAT-box. In *e12*, deletion of Block E eliminated TF binding across the region, indicating that Block E is essential for maintaining local chromatin accessibility. In *e14*, deletion between Block E and the third CCAAT-box repositioned the three CCAAT-boxes in closer proximity, resulting in strong binding at both Block E CCAAT-boxes (1 and 2) and novel binding at the third box (3), indicating enhanced TFs (NF-Ys) recruitment and correlating with elevated *FT* expression and early flowering. In *e16*, removal of both Block E and its immediate downstream sequences placed the third CCAAT-box at the former location of the first CCAAT-box, allowing TFs (NF-Ys) binding and remaining of chromatin accessibility and leading to a wild type-like flowering time.

Our combined genetic and MOA-seq analyses support a model in which the activity of the CCAAT-box module downstream of *FT* is determined not only by motif identity but also by their spatial arrangement and chromatin context (Fig. 4e). In wild type, the first CCAAT/G-box module in Block E is essential and dominant, while the downstream CCAAT-box remains largely inactive, likely constrained by the intervening CArG-containing region and/or an unfavorable chromatin context. In late-flowering *e12* mutants, deletion of Block E CCAAT motifs abolished TF-binding and reduced *FT* expression. By contrast, early-flowering *e13* and *e14* retained Block E but lost the putative repressive barrier, repositioning the downstream CCAAT-box module next to Block E and boosting enhancer activity. In *e16*, deletion of both Block E and the barrier relocated the downstream module to the Block E position, enabling TF binding and restoring wild-type flowering. Together, these findings reveal that the downstream CCAAT-box module acts as a latent enhancer, activated only when placed in a favorable chromatin and spatial context.

## Discussion

Our study uncovers the critical and previously underappreciated role of the 2.3-kb region downstream of *FT*, particularly the distal enhancer Block E, in sustaining full *FT* transcription and ensuring timely flowering under long-day conditions. Genetic deletion of this region, or of key motifs within Block E, caused significant *FT* downregulation and delayed flowering. These findings challenge the historical focus on upstream regulatory sequences and demonstrate that downstream elements can be equally essential. We recommend that future genomic complementation of *ft* mutants, and broader studies of the *FT* regulatory landscape, should include both upstream and downstream sequences to achieve faithful expression with single-copy insertions.

By contrast, deletion of three conserved lncRNAs in the *FT* downstream region had no detectable effect on *FT* expression or flowering time under standard long-day conditions. One of these, *AT1G08757*, shows diurnal expression and hydathode enrichment, features associated with stress or developmental regulation. Although dispensable under our conditions, such lncRNAs may act in specific environmental contexts, such as abiotic stress or pathogen challenge.

Dissection of Block E revealed that the first CCAAT-box and adjacent G-box, separated by only 2 bp, form an essential composite module. This extreme proximity, makes it challenging to selectively disrupt one motif without affecting the other by CRISPR/Cas9, but likely facilitates cooperative action between NF-Y and G-box-binding factors such as PIFs. This interpretation is supported by ChIP-seq studies showing that NF-Y binding sites are frequently enriched not only for CCAAT motifs but also for G-box– containing elements^42^, suggesting NF-Y may also associate with G-boxes either directly or indirectly, potentially through interactions with bZIP or PIF partners. We therefore propose that the first CCAAT-box and adjacent G-box together constitute a core module within Block E enabling NF-Y–mediated transcriptional activation of *FT*.

Not all conserved motifs in accessible chromatin were essential. The second CCAAT-box, RE-α, and a TBS-like motif within Block E made modest or context-dependent contributions. Their effects often differed between developmental stages or time points, consistent with conditional activity. RE-α deletion transiently increased *FT* expression early in development, in line with its proposed role in repression during darkness^43^. Instead of promoting flowering, the TBS-like motif deletion caused mild late flowering, suggesting it contributes to balancing repressor activity rather than acting as a strict silencer.

A downstream CCAAT-box located beyond Block E was largely inactive in wild type, showing little TF binding by MOA-seq. This inactivity is likely due to an unfavorable chromatin context or to interference from the intervening CArG-box containing region, which may recruit repressors or influence nucleosome positioning. Strikingly, deletion of this barrier in *e13*/*e14* repositioned the downstream module adjacent to Block E, enabling TF binding and elevated *FT* expression. In *e16*, relocation of this module to the Block E position supported near wild-type flowering. Thus, the downstream CCAAT-box can function as a latent enhancer when placed in a favorable spatial and chromatin context. Analogous to findings in *Drosophila* where Zelda- or Grainyhead-biased sequences can prime genomic regions to act or evolve as enhancers, our results suggest that the CCAAT-box represents a plant-specific manifestation of this general principle^44^.

Together, these findings highlight a modular regulatory architecture in which enhancer activity depends not only on motif identity but also on spatial arrangement and chromatin context. This modularity provides a mechanistic explanation for how regulatory sequences can buffer genetic variation: critical motifs in dominant modules (e.g., Block E) ensure robust *FT* induction, whereas downstream latent modules can compensate under altered genomic contexts. More broadly, plant *cis*-regulatory elements may frequently harbor latent enhancer activity that is constrained in their native genomic contexts. Such latent modules could serve as evolutionary “reserve” elements, capable of being activated by sequence rearrangements, deletions, or chromatin remodeling, potentially contributing to phenotypic plasticity and adaptive evolution. Future work will be needed to determine how widespread such context-dependent enhancer activation is across plant genomes and whether similar principles govern other developmental regulators.

In conclusion, the *FT* downstream regulatory landscape is highly modular and position-sensitive, integrating multiple activators and repressors within a dynamic chromatin environment. This architecture may enable evolutionary flexibility in flowering-time control while preserving core regulatory logic. The principles uncovered here— particularly the positional activation of latent enhancer modules—may inform strategies to engineer precise gene expression in diverse plant species and potentially beyond the plant kingdom.

## Methods

### Plant material and growth conditions

The Columbia (Col-0) ecotype of *Arabidopsis thaliana* was used as the wild type and for CRISPR/Cas9 mutagenesis. The *ft-10* mutant was described previously^45^. For flowering time analysis, seeds were stratified in the dark at 4 °C for two days and then grown on soil in a greenhouse or a growth chamber under long-day conditions (16 h light/8 h dark, 20–24 °C). For gene expression analysis and MOA-seq, surface-sterilized seeds were sown on MS medium (4.4 g MS salts, 0.5 g MES, 10 g sucrose, 9 g PhytoAgar per liter, pH 5.7). After two days of stratification at 4 °C, plates were transferred to a growth chamber (Percival) under long-day conditions (16 h light/8 h dark, 120–150 µmol m⁻² s⁻¹, 21 °C).

### Comparative Genomic Analysis Using VISTA-Point

To identify conserved regions downstream of *FT* across Arabidopsis accessions and relative species, the VISTA-Point (http://genome.lbl.gov/vista/) was used with pre-computed genome datasets available in its database. The *A. thaliana* genome was selected as the base genome. Conservation thresholds were set to 70% sequence identity across a 100 bp window. The tool visualized sequence conservation within and downstream the *FT* locus, with a focus on non-coding sequences to infer potential regulatory elements.

### Generation of mutants by CRISPR/Cas9 technology

CRISPR/Cas9 constructs were generated using the genome editing toolkit as described previously^46^. In brief, sgRNAs were designed with CRISPR-P 2.0^47^, annealed oligonucleotides were assembled into shuttle vectors via simultaneous restriction/ligation with *BbsI-HF* (NEB), and subsequently cloned into the dicot genome-editing recipient vector pDGE347 by Golden Gate assembly with *BsaI-HF* (NEB). The resulting CRISPR/Cas9 constructs, carrying sgRNAs targeting the *FT* downstream region, were transformed into Agrobacterium strain GV3010(*p*Soup) and introduced into Col-0 plants by floral dip in Agrobacterium suspension (5% sucrose, 0.02% Silwet L-77)^48^.

Large-deletion mutants (*d3*, *d4*, *d5*) and the small-deletion mutant (*e7*) were generated by crossing primary T1 transformants with wild type, following a previously described strategy^49^. Indels were detected in F1 plants by PCR using primers flanking the target sites and confirmed by Sanger sequencing. CRISPR/Cas9-free homozygous mutants were obtained in the F2 or F3 generation and used for further experiments. For other mutants, indels were identified in T1 or T2 plants, and CRISPR/Cas9-free homozygous lines were selected in the T3 or T4 generation. PCR primers and sgRNA oligonucleotides were synthesized at SIGMA listed in Supplementary Table 1.

### GUS reporter construct and staining

To generate the binary vector *AT1G08757p-GUS*, a 1.5-kb promoter fragment together with 222 bp of the *AT1G08757* were amplified from Col-0 genomic DNA and cloned into the previously developed *GW-GUS-pGREEN* vector^4^ using NotI and HindIII restriction sites. Primer sequences used for amplification are listed in Supplementary Table 1. GUS staining was performed as previously described^4^.

### Generation of complementation constructs and transgenic plants

A 12.6 kb *FT* genomic fragment (spanning 8.1 kb upstream of the start codon to 2.3 kb downstream of the stop codon) was subcloned from the BAC clone F5I14 into the *pGAP-Km* vector by homologous recombination as described previously^50^. To enable homologous recombination, two ∼600 bp PCR fragments (FLANK1 and FLANK2) were amplified and inserted into *pGAP-Km*: FLANK1 cleaved with *BamHI* and *SalI*, FLANK2 cleaved with *SalI* and *EcoRI*, and the vector cleaved with *BamHI* and *EcoRI*. The resulting FLANK1–FLANK2–pGAP-Km construct was introduced into *E. coli* SW102 cells harboring BAC F5I14, and recombination was induced. PCR primers used for FLANK amplification are listed in Supplementary Table 1.

The final construct, *pGAP-12.6kFT*, was transformed into Agrobacterium strain GV3010(*p*MP90RK) and introduced into *ft-10* plants by floral dipping^48^.

### Copy number testing

A plasmid containing fragments of *PP2A* and *KanR* cloned in single copy into a pBluescript KSII backbone was diluted to generate standard curves for absolute quantification of qPCR reactions. The ratios between *KanR* and *PP2A* signals were determined for standard reactions and for those using DNA prepared from selected transgenic T2 or T3 lines as template. Primers are listed in Supplementary Table 1.

### Flowering time analysis

Flowering time was assessed by counting the total number of rosette and cauline leaves (including cotyledons) at bolting. Leaf numbers were averaged from at least 11 plants per genotype.

### Total RNA extraction and gene expression analysis

Seedlings (12–14 days old) grown under long-day conditions were harvested either every 4 hours from ZT0 to ZT24 or at defined time points (ZT4 and/or ZT16), as described in the figure legends. Total RNA was extracted using TRIzol™ Reagent (Ambion, Life Technologies), and genomic DNA was removed with the DNA-free™ DNA Removal Kit (Invitrogen, AM1906).

For temporal expression profiling of *FT* and non-coding RNAs, one-step RT-qPCR was conducted with 500 ng of DNA-free RNA in a 10 µL reaction using the iTaq^TM^ Universal SYBR^®^ Green One-Step Kit (Bio-Rad) on a CFX384 Real-Time System (Bio-Rad). For comparative analysis of *FT* expression in wild type and mutants, 2 µg of DNA-free RNA was reverse transcribed into cDNA in a 20 µL reaction using oligo(dT) primers and SuperScript IV Reverse Transcriptase (Invitrogen, Thermo Fisher Scientific). Quantitative PCR was then performed with 1 µL of cDNA in a 10 µL reaction using iQ^TM^ SYBR^®^ Green Supermix (Bio-Rad) on the CFX384 Real-Time System (Bio-Rad), with PP2A as an internal control for normalization. Primers are listed in Supplementary Table 1.

### MOA-seq and data analysis

MOA-seq was performed as described previously^51^, with minor modifications. Briefly, ∼80 mg of frozen, N_2_-ground 12-day-old seedlings were fixed in a freshly prepared solution of 1% paraformaldehyde in a fixation buffer (18 mM PIPES, 80 mM KCl, 20 mM NaCl, 2 mM EDTA, 0.5 mM EGTA, 384 mM sorbitol, 1.2 mM DTT, 0.18 mM spermine, 0.6 mM spermidine, 0.2 mM phenanthroline, 0.4 mM PMSF, and 2 µg/ml each of aprotinin and pepstatin A) for 10 mins, quenched with glycine, and nuclei were isolated in MDB buffer (50 mM HEPES, 12.5% glycerol, 25 mM KCl, 4 mM MgCl₂, 1 mM CaCl₂, pH 7.6) by homogenization and filtration. Nuclei were resuspended in 70 μl MDB supplemented with 1% Triton X-100 and digested with MNase (50 or 100 U/ml; buffer only for control) at 37 °C for 15 min. Reactions were stopped with EGTA and RNase A, followed by Proteinase K decrosslinking at 65 °C overnight. DNA was purified using the NEB Monarch PCR & DNA Cleanup Kit, quantified with the Qubit dsDNA HS Assay Kit, and checked on agarose gels.

Libraries were prepared with the NEBNext® Ultra II DNA Library Prep Kit (Illumina®) following the same modifications as Engelhorn et al^51^. Sequencing was performed on an Illumina NovaSeq 6000 S4 (WT Rep1, *e16* Rep1) or on the DNBSEQ G400/PE85 platform (BGI; all other replicates) in paired-end mode.

Adapters were trimmed with SeqPurge^52^ using default settings, ensuring a minimum read length of 20 bp. Paired-end reads were merged with FLASH^53^. Reads were aligned to the *A. thaliana* TAIR10 genome using STAR^54^, with maximum permissible intron size restricted to 1 (--alignIntronMax 1) and multimapping tolerance set to 1 (--outFilterMultimapNmax), allowing only uniquely mapping reads. Alignments were converted to coordinate-sorted BAM files and coverage tracks generated with deepTools bamCoverage^55^ using CPM normalization, 1-bp bins, maximum fragment length of 80 bp, and minimum mapping quality of 255. Coverage was visualized using IGV^56^.

### Statistical analyses

Statistical tests were performed as described in the figure legends. Pairwise comparisons were made using Welch’s *t*-test (ns p>0.05, * p<0.05, ** p<0.01, *** p<0.001, **** p<0.0001). Multiple-sample comparisons were determined by one-way ANOVA followed with post-hoc Tukey HSD, and significant differences at the *P* < 0.01 level are indicated by different lowercase letters.

## Acknowledgments

We thank the Jane E. Parker group for sharing shuttle and recipient vectors, originally developed by the Johannes Stuttmann group. We also thank the Csaba Koncz group for pGAP-pBR-Km and the Christian Fankhauser group for sharing the *pif457* triple mutant. FT and HRZ are supported by a Core Grant from the Max Planck Society. FT also receives funding from the DFG through the Cluster of Excellence CEPLAS (EXC 2048/1 Project ID: 390686111).

## Data availability

The raw MOA-seq data described in this paper have been deposited at NCBI SRA under accession number PRJEB97763.

## Extended Data

**Extended Data Fig. 1:**
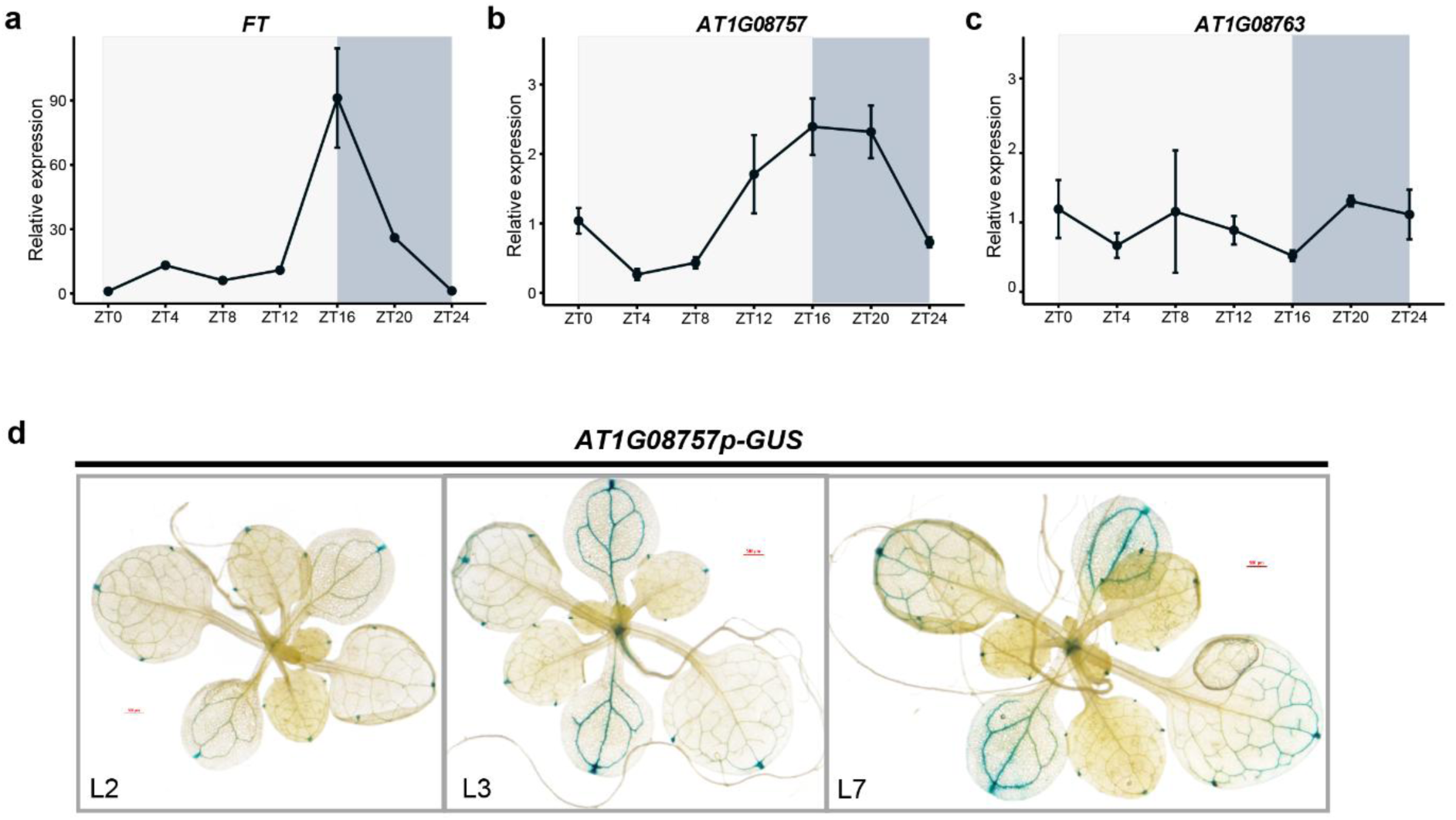
Spatiotemporal expression pattern of the non-coding gene *AT1G08757*. **a–c**, Transcript accumulation measured by one step RT-qPCR from 14-day old seedlings grown in under long-day (LD) at 21°C in condition. Samples were collected in 4h intervals over a time-period of 24h. Error bars show standard error of the mean of three biological replicates. (**a**) *FT*, (**b**) non-coding transcript *AT1G08757*, (**c**) non-coding transcript *AT1G08763*. **d**, GUS staining of three independent *AT1G08757p*-*GUS* transgenic lines(L) grown under LD conditions. Scale bar: 500 µm.

**Extended Data Fig. 2:**
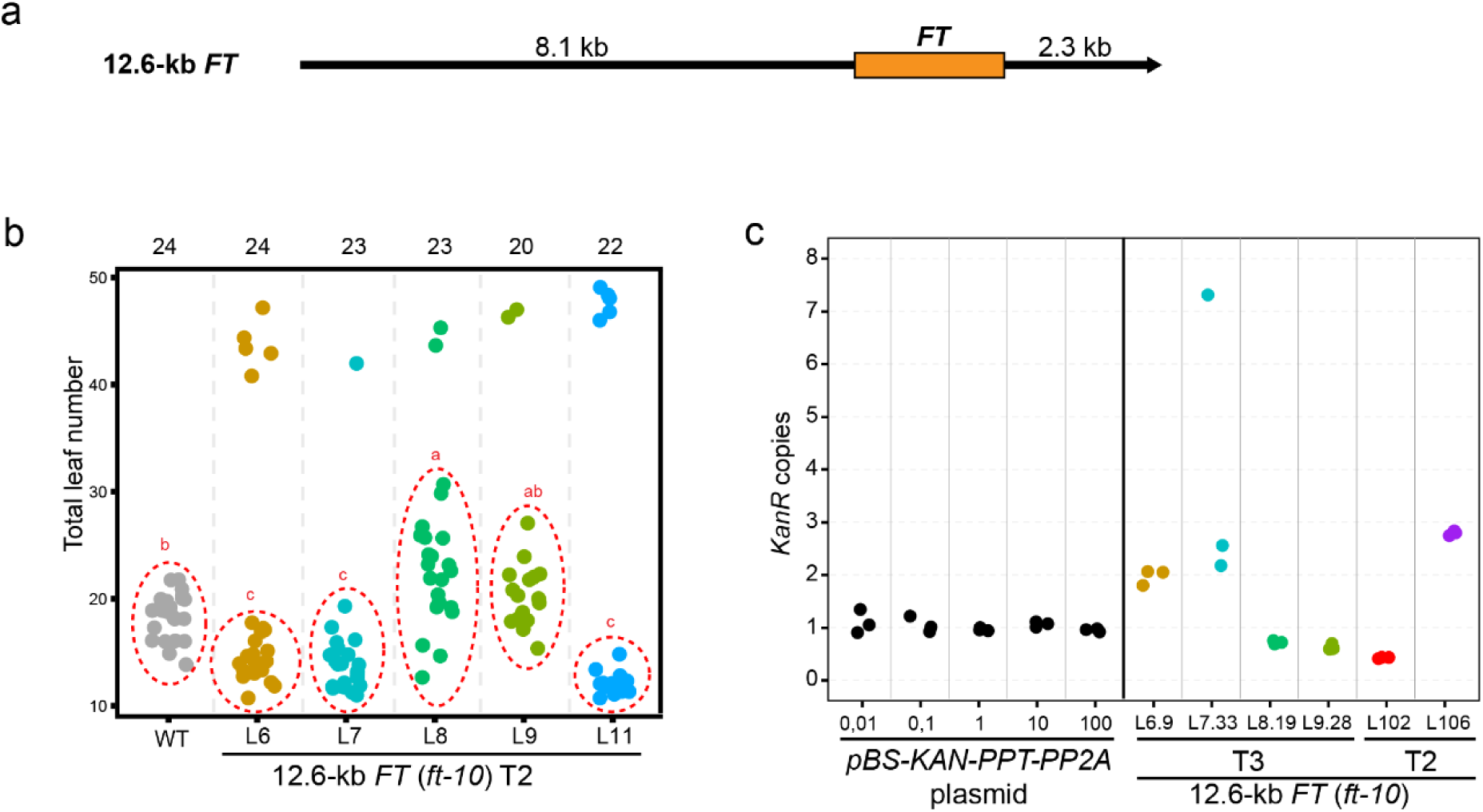
Full complementation of the late-flowering *ft-10* phenotype by *FT* with upstream and downstream regulatory elements. **a**, Schematic representation of *FT* and its flanking sequences retrieved from BAC clone F5I14 by recombineering. The 12.6-kb *FT* genomic fragment (Chr1:24323415– 24335986) spans 8.1 kb upstream of the start codon through 2.3 kb downstream of the stop codon, encompassing all known regulatory elements. **b**, Flowering time, measured as the number of leaves at bolting, in WT and T2 populations of five independent transgenic lines carrying the 12.6-Kb *FT* genomic region. Replicate numbers are shown above the plot. For each line, plants flowering similarly to WT (dashed circles) were analyzed with WT (also circled) by ANOVA with Tukey’s HSD. Different letters indicate significant differences. **c**, Copy-number test of transgenic lines by qPCR. The ratio of *KanR*/*PP2A* determined from a reference plasmid (left panel) was compared to values obtained from T3 offspring of lines shown in b and from T2 plants of two additional lines. Values show technical replicates of qPCR results. Lines L8, L9 and L102 were evaluated as single copy, note that lines L7, L102, and L106 were not homozygous.

**Extended Data Fig. 3:**
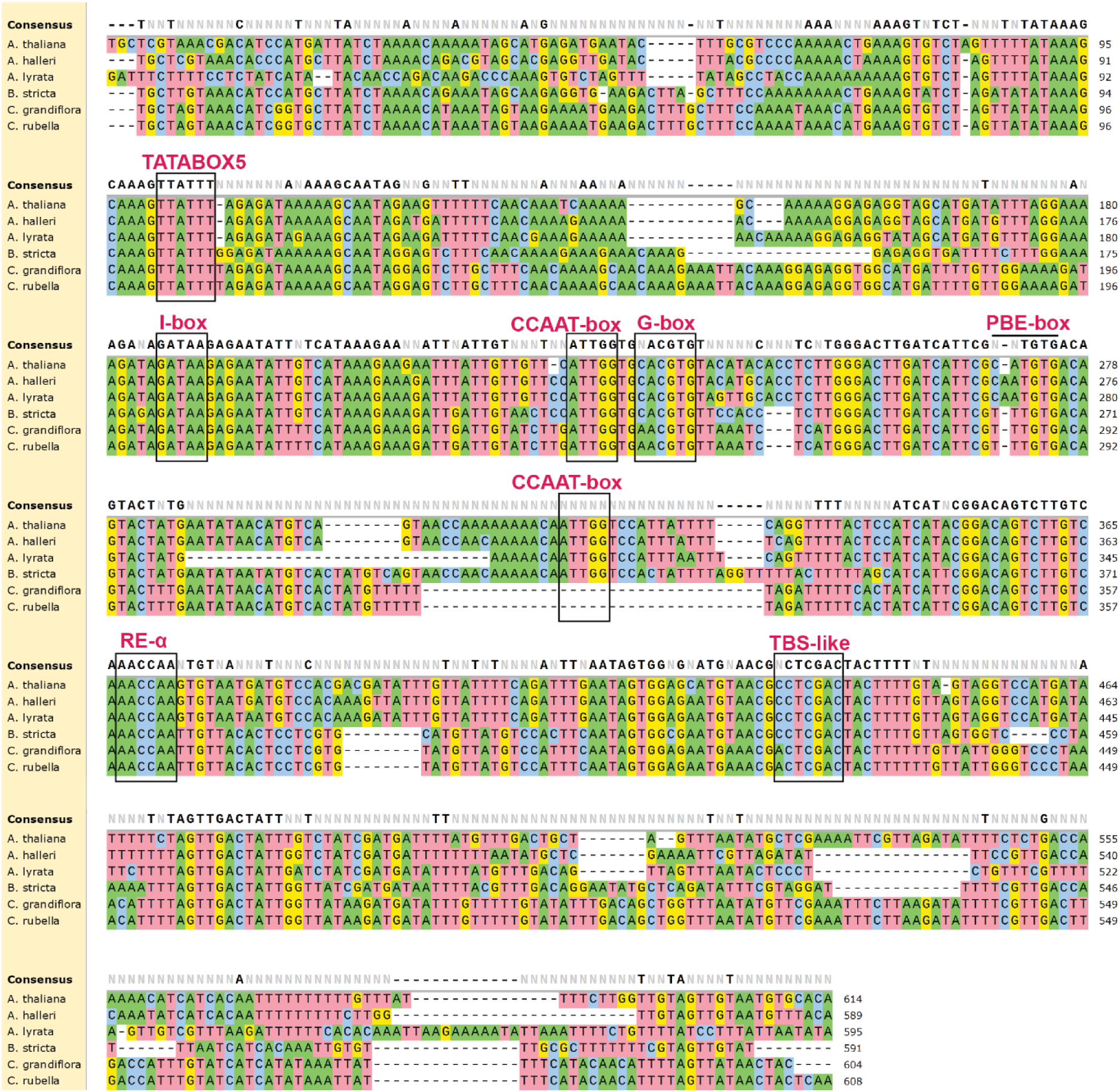
Conservation of *cis*-regulatory DNA motifs within Block E across Brassicaceae species. The motifs TATABOX5, I-box, CCAAT-box, G-box, PBE-box, and RE-alpha were identified within *A. thaliana* Block E (Chr1:24,335,009–24,335,622) using the New PLACE database (https://www.dna.affrc.go.jp/PLACE/?action=newplace)^57^. The TBS-like motif, also known as MA0553.1, was previously described^20,32^. Sequence alignment of Block E was performed using Clustal Omega in SnapGene across six related species: *A. thaliana*, *A. halleri*, *A. lyrata*, *B. stricta*, *C. grandiflora*, and *C. rubella*. Conserved occurrences of the motifs are highlighted with black boxes.

**Extended Data Fig. 4:**
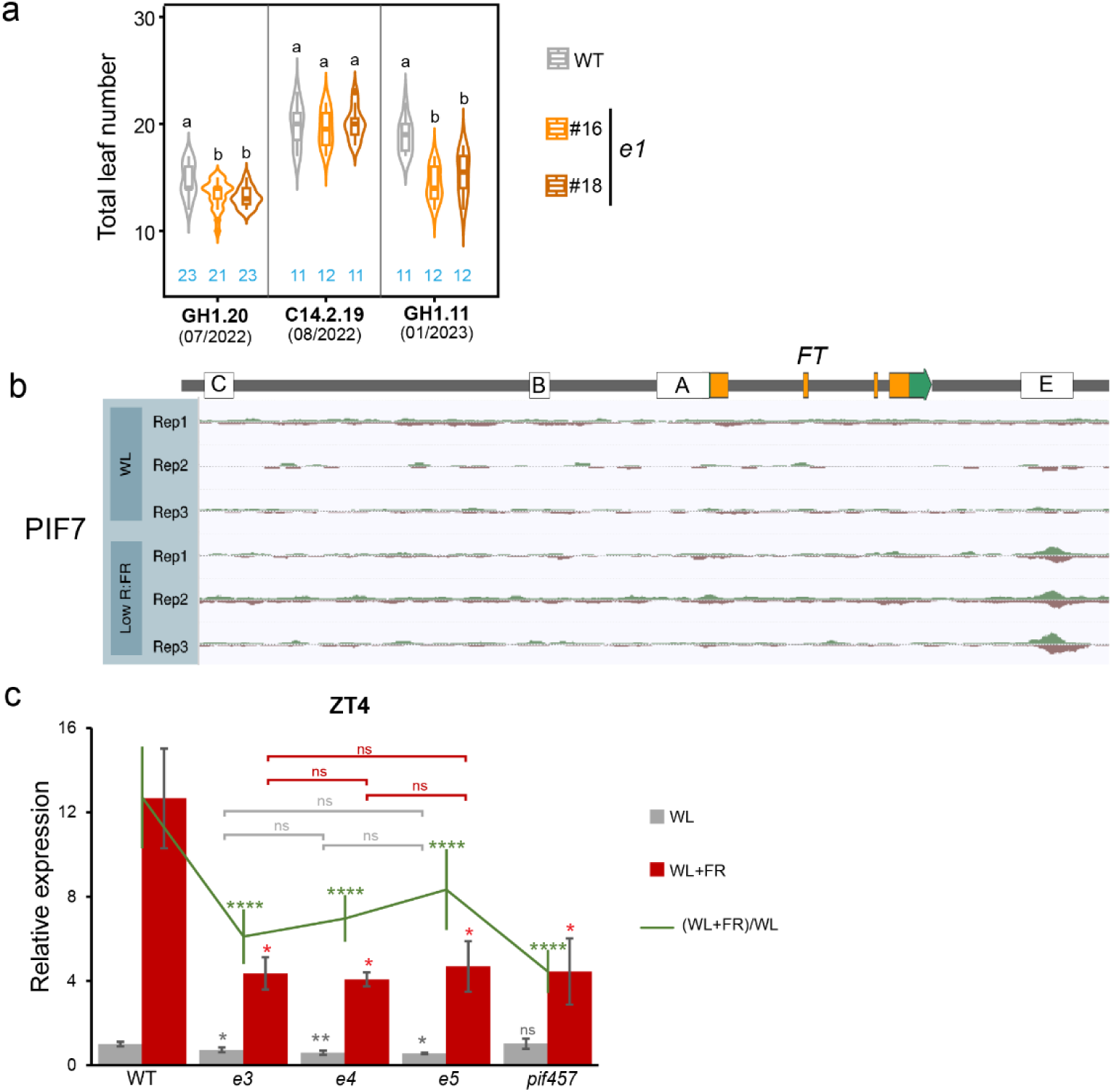
Neither the upstream G-box nor the G-box alone in Block E could compensate the loss of CCAAT-box and G-box within Block E. **a**, Flowering time measured as leaves produced at the main shoot until flowering in wild type (WT) and two independent segregants of a deletion after the second G-box downstream of *FT* (*e1*). Plants were grown in either greenhouse (**GH**) or growth chamber (**C**) conditions. The number of replicates is indicated in blue, statistical test by ANOVA with posthoc HSD Tukey testing. Statistically significant differences are indicated by different letters. **b**, Schemes of the *FT* locus including its 5.7 kb promoter and 2 kb downstream region, with Blocks A, B, C, and E indicated. A snapshot of the PIF7 ChIP-seq data around the *FT* locus is also shown, which is available at (http://neomorph.salk.edu/aj2/pages/hchen/)^34^. **c**, *FT* transcript accumulation was measured by RT-qPCR in the indicated genotypes. Samples collected at ZT4 from 12-day old seedlings grown in long days (LD) with white light (WL) and from seedlings grown 7 days in LD (WL) followed by 5 days with additional far-red light (WL+FR). The red-to-far-red light ratio was 3.3 under WL and 0.27 under WL+FR. Analysis was using the 2deltaCq method using Col-0 as reference and *PP2A* as housekeeping control. Error bars show standard error of the mean (SEM) of three biological replicates. The green line shows the ratio of *FT* transcript accumulation between WL+FR and WL for each genotype, with SEM calculated from nine ratios across three biological replicates per condition. Statistical analysis by Welch’s *t*-test, significant differences indicated by stars (ns p>0.05, * p<0.05, ** p<0.01, *** p<0.001, **** p<0.0001). Symbols (or ns) above bars or the line denote significance relative to WT, with colors indicating the corresponding conditions. Bracketed ns symbols indicate comparisons among Block E mutants, with colors specifying conditions.

**Extended Data Fig. 5:**
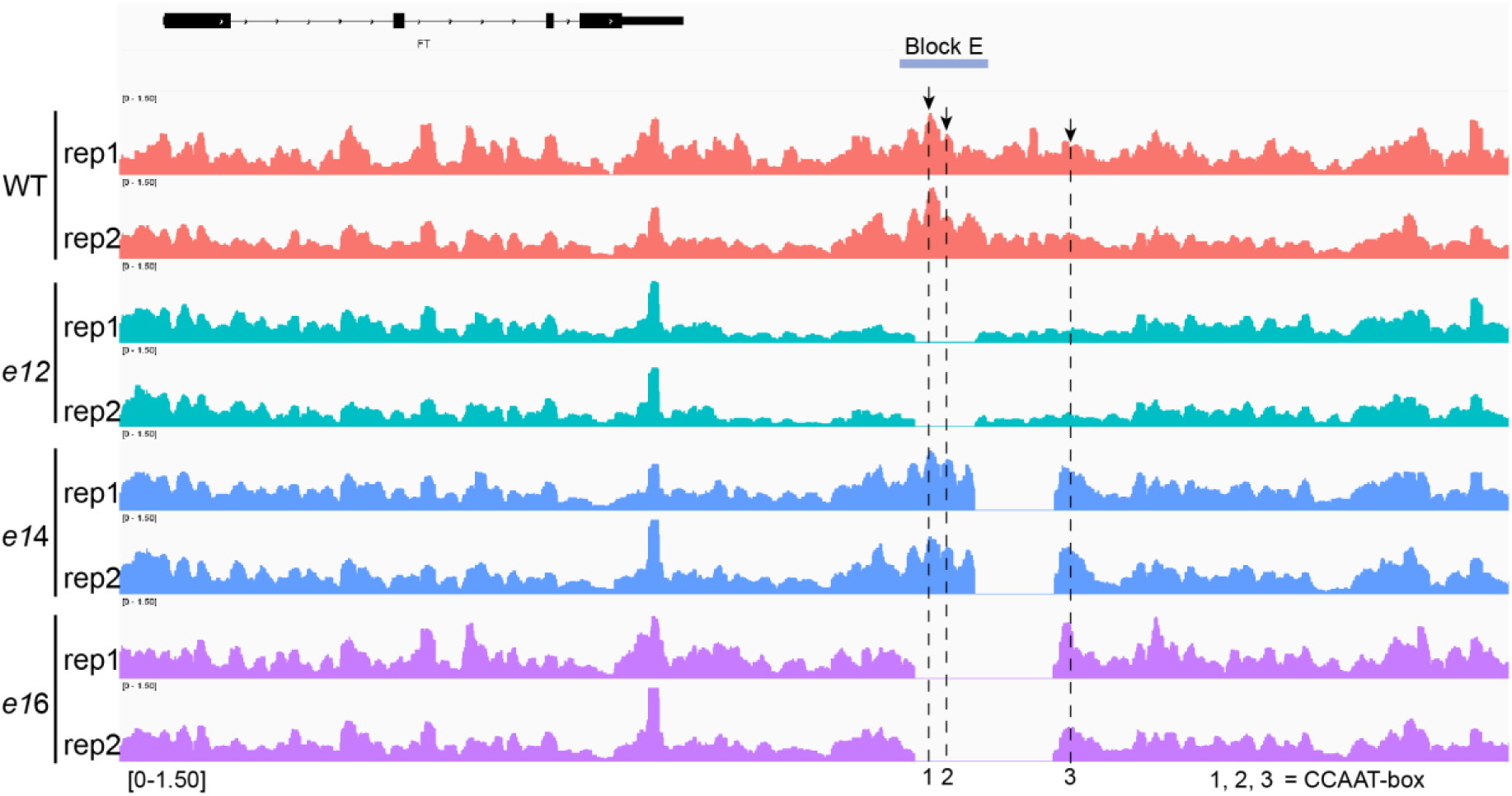
Altered transcription factor occupancy at Block E and downstream CCAAT-boxes in Block E deletion mutants. MOA-seq read coverage from two biological replicates (rep1 and rep2) at the *FT* locus and downstream region in wild type (WT) (red), *e12* (cyan), *e14* (blue), and *e16* (purple). The *FT* gene structure is shown above (black boxes, exons and UTRs; black lines, introns). Block E is indicated by a light blue line. Numbers 1–3 denote the three CCAAT-box motifs within Block E and its downstream region; their exact positions are indicated by black arrows and corresponding dashed lines.

**Supplementary Table 1:**
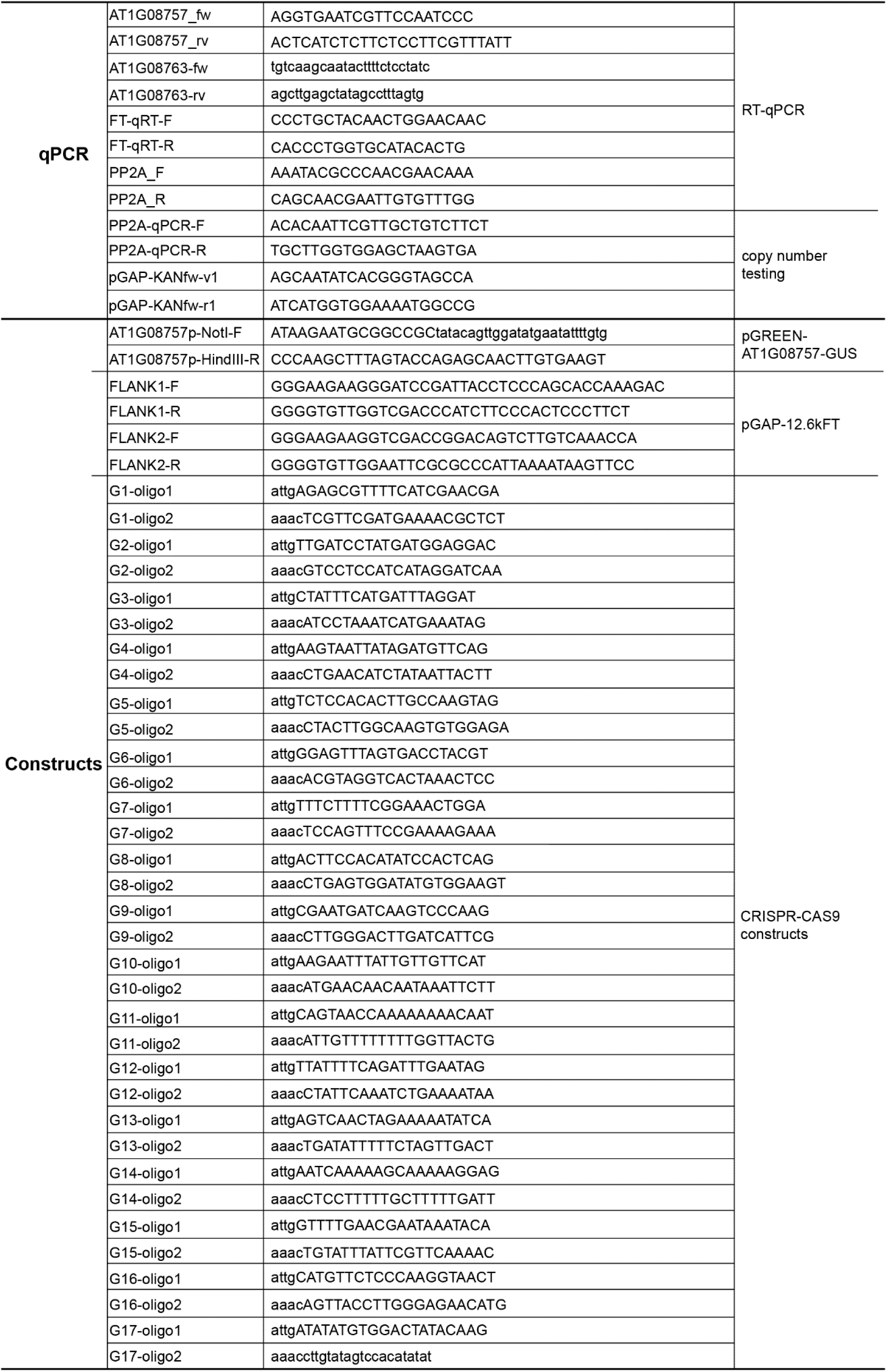
Oligonucleotide sequences used in this study.

## Notes

### Competing Interest Statement

The authors have declared no competing interest.

